# Rapid parallel adaptation in distinct invasions of *Ambrosia artemisiifolia* is driven by large-effect structural variants

**DOI:** 10.1101/2024.09.18.613801

**Authors:** Paul Battlay, Samuel Craig, Andhika R. Putra, Keyne Monro, Nissanka P De Silva, Jonathan Wilson, Vanessa C. Bieker, Saila Kabir, Nawar Shamaya, Lotte van Boheemen, Loren H. Rieseberg, John R. Stinchcombe, Alexandre Fournier-Level, Michael D. Martin, Kathryn A. Hodgins

## Abstract

When introduced to multiple distinct ranges, invasive species provide a compelling natural experiment for understanding the repeatability of adaptation. *Ambrosia artemisiifolia* is an invasive, noxious weed and chief cause of hay fever. Leveraging over 400 whole-genome sequences spanning the native range in North America and two invasions in Europe and Australia, we inferred demographically distinct invasion histories on each continent. Despite substantial differences in genetic source and effective population size changes during introduction, scans of both local climate adaptation and divergence from the native range revealed genomic signatures of parallel adaptation between invasions. Disproportionately represented among these parallel signatures are 37 large haploblocks – indicators of structural variation – that cover almost 20% of the genome and exist as standing genetic variation in the native range. Many of these haploblocks are associated with traits important for adaptation to local climate, like size and the timing of flowering, and have rapidly reformed native range clines in invasive ranges. Others show extreme frequency divergence between ranges, consistent with a response to divergent selection on different continents. Our results demonstrate the key role of large-effect standing variants in rapid adaptation during range expansion, a pattern that is robust to diverse invasion histories.

## Introduction

Invasive species are one of the key drivers of ecological change globally (Arneth et al., 2019; Pejchar and Mooney, 2010), posing a significant and growing threat to global biodiversity, agriculture and human health. They are a leading cause of species extinction worldwide (Clavero and García-Berthou, 2005; Pimentel et al., 2005), and the global cost of invasive species – predominantly agricultural yield loss and control measures – was estimated to exceed 423 billion US dollars for 2019 alone (Fokam et al., 2023). Human intervention has so far been insufficient to control the proliferation of biological invasions (Seebens et al., 2017), which are predicted to accelerate as climate change increases (Dukes and Mooney, 1999), pointing to the need for a greater understanding of the mechanisms governing invasion success. Here we use population genomics to determine the genetic mechanisms underlying invasion success in a prolifically invasive weed.

A long-standing mystery in invasion biology is how invasive species can be so successful despite founder effects and bottlenecks that are expected during initial colonization (i.e., the genetic paradox of invasion; Allendorf and Lundquist, 2003; Estoup et al., 2016; Schrieber and Lachmuth, 2017). Such demographic changes should reduce genetic variation (Willi et al., 2006), cause inbreeding (Angeloni et al., 2011; Fauvergue et al., 2012), and increase the frequency of deleterious alleles (Willi et al., 2013), resulting in depressed fitness and limited adaptive potential. This paradox may be resolved in some cases by repeated introductions which can lead to high levels of genetic variation in introduced populations thereby mitigating these expected fitness costs associated with colonization (Bieker et al., 2022; Dlugosch et al., 2015; Dlugosch and Parker, 2008; McGoey and Stinchcombe, 2021). However, few studies have leveraged invasions with contrasting demographic histories within a single species to assess its influence on the evolutionary trajectories of populations.

When introduced to multiple distinct ranges, invasive species provide a compelling natural experiment for understanding the repeatability in the genetic basis of adaptation. In contrast to the infinitesimal architecture theorized to underlie most adaptive traits (Fisher, 1919, 1930), mutations with larger effects on traits are more likely to contribute to adaptation under certain circumstances. For example, when there has been a sudden, large shift in adaptive optima (Orr, 1998), when populations are small (Charlesworth, 2009) or declining (McDonough and Connallon, 2023), or if selective pressures vary across space resulting in local adaptation (Yeaman and Whitlock, 2011). Large-effect genetic architectures are more likely to result in repeated use of the same gene for adaptation (Yeaman et al., 2018), particularly when adaptation occurs from standing variation (Ralph and Coop, 2015), and when there is gene flow between locally adapted populations (Battlay, Yeaman, et al., 2024). Consistent with theoretical predictions, large structural variants are increasingly being identified as drivers of rapid adaptation in invasive species (Battlay et al., 2023; Battlay, Hendrickson, et al., 2024; Ma et al., 2024; Santangelo et al., 2022; Tepolt and Palumbi, 2020; Wilson et al., 2024). However, there have been only limited empirical studies of parallelism between invasions at the genomic level (but see Battlay, Hendrickson, et al., 2024; Olazcuaga et al., 2020; van Boheemen and Hodgins, 2020). Moreover, different invasions may be subjected to population bottlenecks of varying severity, and sourced from distinct genetic clusters in the native range or other invasive ranges, both of which would reduce genomic parallelism. It is unclear to what extent such demographic factors influence parallelism, and hence the predictability, of rapid adaptation.

*Ambrosia artemisiifolia* is a noxious weed, and its airborne pollen is a chief cause of hay fever (allergic rhinitis; Caillaud et al., 2014; Della Torre et al., 1996). As a result of human introductions over the last 200 years, this North American native has successfully colonized all continents except Antarctica. Its recent, aggressive expansion across the globe makes the species a powerful model for understanding parallel rapid adaptation. The reemergence of latitudinal clines in traits in invasive *A. artemisiifolia* populations suggests the involvement of rapid adaptation in the species’ spread (Li et al., 2015; McGoey et al., 2020; van Boheemen et al., 2019), while genomic data have highlighted the role of large-effect standing variants in the successful invasion of Europe (Battlay et al., 2023). Here we sequenced 95 whole genomes spanning *A. artemisiifolia’s* invasive Australian range. When combined with 348 whole-genome sequences of contemporary samples from North America and Europe (Bieker et al., 2022), we were able to characterize the invasion history and contrast adaptive change during invasions on two continents. We reconstructed genetically and demographically distinct invasion histories in each introduced range with evidence of a substantial bottleneck in Australia but not in Europe. We then tested the hypothesis that genetic bottlenecks should limit the extent of parallel adaptive evolution from standing variation by comparing the extent of parallel signatures of adaptation between the introduced ranges and discovered that genomic repeatability was high regardless of the invasion history. This repeatability occurred disproportionately in large structural variants that cover ∼20% of the genome and underpin substantial variation in locally adaptive traits such as flowering time. These results underscore the importance of standing variation in large effect structural variants in rapid, repeated adaptation during range expansions with distinct demographic histories.

## Results

### Contrasting invasion histories in Europe and Australia

A principal component analysis (PCA) of genetic variation across 443 whole genome-sequenced *A. artemisiifolia* samples (Fig. 1A; Table S1) revealed little evidence of divergence between North America and Europe, consistent with previous inferences of repeated introductions from North America (Bieker et al., 2022; van Boheemen et al., 2017; Fig. 1B). Most European samples overlapped with North American mid east and west clusters, with no evidence that the North American south cluster (which comprises samples from Florida) contributed to European introductions (Fig. 1B). In contrast to the genetic similarity between North American and European samples, there is substantial divergence between the main genetic cluster of Australian samples and samples from the other ranges. The main Australian cluster shows the highest similarity with the North American south cluster (Fig. 1B), but Australian samples outside this main cluster group more closely with North American east samples (Fig. 1B) and are largely found in the southernmost Australian population (Nelligen, NSW; Fig S1). Admixture proportions measured across samples support these observations (Fig. S2). These results suggest European and Australian invasive populations were founded by distinct sources within the native range.

**Figure 1.**
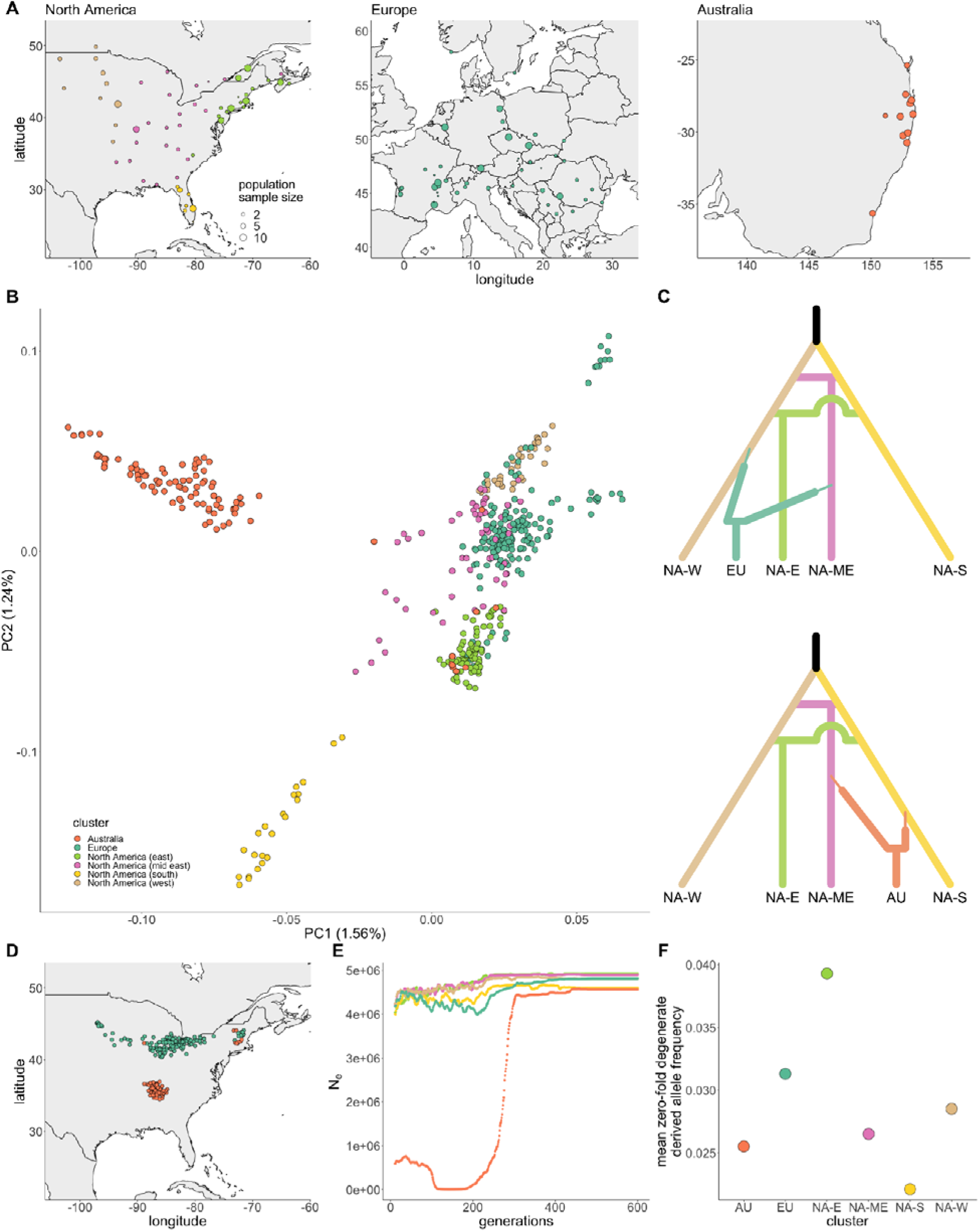
Invasion sources and recent demography of *A. artemisiifolia*. **A.** Sampling locations for 443 *A. artemisiifolia* whole-genome sequences spanning 119 populations across three ranges. **B.** Mapping of individual samples along the first two principal components of 100,000 linkage disequilibrium-thinned genetic variants sampled from outside genes and haploblocks, coloured by North American cluster or invasive range. **C.** The most likely scenarios from DIY-ABC modeling historic divergence between native-range genetic clusters (NA-W: North America west; NA-ME: North America mid east; NA-S: North America south; NA-E: North America east) and introductions to Europe (EU) and Australia (AU). Time is not to scale. **D.** Deep neural network-predicted source locations in North America for invasive European and Australian samples. **E.** Effective population sizes (N_e_) over time inferred from linkage disequilibrium in each North American cluster or invasive range. **F.** Mean allele frequency at zero-fold degenerate sites for invasive ranges and North American clusters. Based on 95% CIs from bootstrapping sites, all points are significantly different from one another. CIs are too narrow to show on the plot.

To determine the most likely scenarios of divergence and admixture in the four native range genetic-spatial clusters (Bieker et al., 2022) and the two invasive ranges, we used *DIYABC-RF* (Collin et al., 2021). The best-supported scenario for the development of genetic structure in the native range modeled the division of a common ancestral population into south and west clusters, with the independent admixture of these clusters resulting in the formation of the mid east and east clusters (posterior probability *p* = 0.534; Fig. S3; Table S2). The best-supported scenario for the European invasion modelled a primary introduction from the west cluster followed by an independent secondary introduction from the mid east cluster, whereas the best-supported Australian scenario modelled introductions from the mid east cluster and south cluster with ambiguity regarding order of arrival (posterior probability *p* = 0.473; Fig. 1C; Fig. S4; Table S3; Table S14).

To further identify the invasion sources, we trained a deep neural network on genetic variation from the North American range using *Locator* (Battey et al., 2020), and predicted source locations for samples from each invasive range. *Locator* results were consistent with those from PCA and *DIYABC-RF*, with most European samples showing northern North American ancestry and most Australian samples showed southern North American ancestry (Fig. 1D). The Australian samples that grouped separately from the main Australian cluster on the PCA were predicted to have been sourced from much further to the north and east of North America (Fig. S2), consistent with their location on the PCA. Comparisons of climatic data for introductions and their putative sources revealed patterns of preadaptation. Australian populations share a similar climate niche with their putative source populations in the North America south cluster, while European populations are more climatically similar to the more northern parts of the North American range (Fig. S5). This is reiterated by the strong relationship between absolute population latitude and *Locator* source location prediction (Fig. S5).

Linkage disequilibrium-based estimates of effective population size using *GONE* (Novo et al., 2023) in each range showed evidence of a recent population bottleneck in Australia (a 416-fold reduction in effective population size) but not in North America or Europe (Fig. 1E). We also measured the average frequency of derived zero-fold degenerate alleles – a metric designed to estimate genetic load (Simons and Sella, 2016) – in North American clusters and each invasive range. Load estimates in each invasive range reflected their North American cluster ancestry rather than demographic events during invasion (Fig. 1F). Furthermore, similar patterns were observed in putatively neutral four-fold degenerate sites (Fig. S6). Together these results suggest the invasions of Europe and Australia were distinct both in terms of genetic sources and demographies.

### Large structural variants support adaptive trait evolution

By analyzing local genomic population structure across all 443 samples in our dataset, we identified and genotyped 37 haploblocks (population-genomic signatures of large structural variants; Table S4), including the 15 previously described in (Battlay et al., 2023). Haploblocks ranged in size from 0.7 Mbp to 17.3 Mbp, cumulatively covered 18.5% of the genome, and varied in frequency between ranges (Fig. 2A). Fifteen of the 37 haploblocks (40%) corresponded to structural variants that were heterozygous in our diploid reference assembly (Fig. S6; Table S4). While all haploblocks heterozygous in the reference include inversions, some are more complex (e.g., *hb-chr9* includes a large deletion in the inverted haplotype, while *hb-chr8a* consists of three tandem inverted regions that appear to segregate as a single structural variant; Fig. S6).

**Figure 2.**
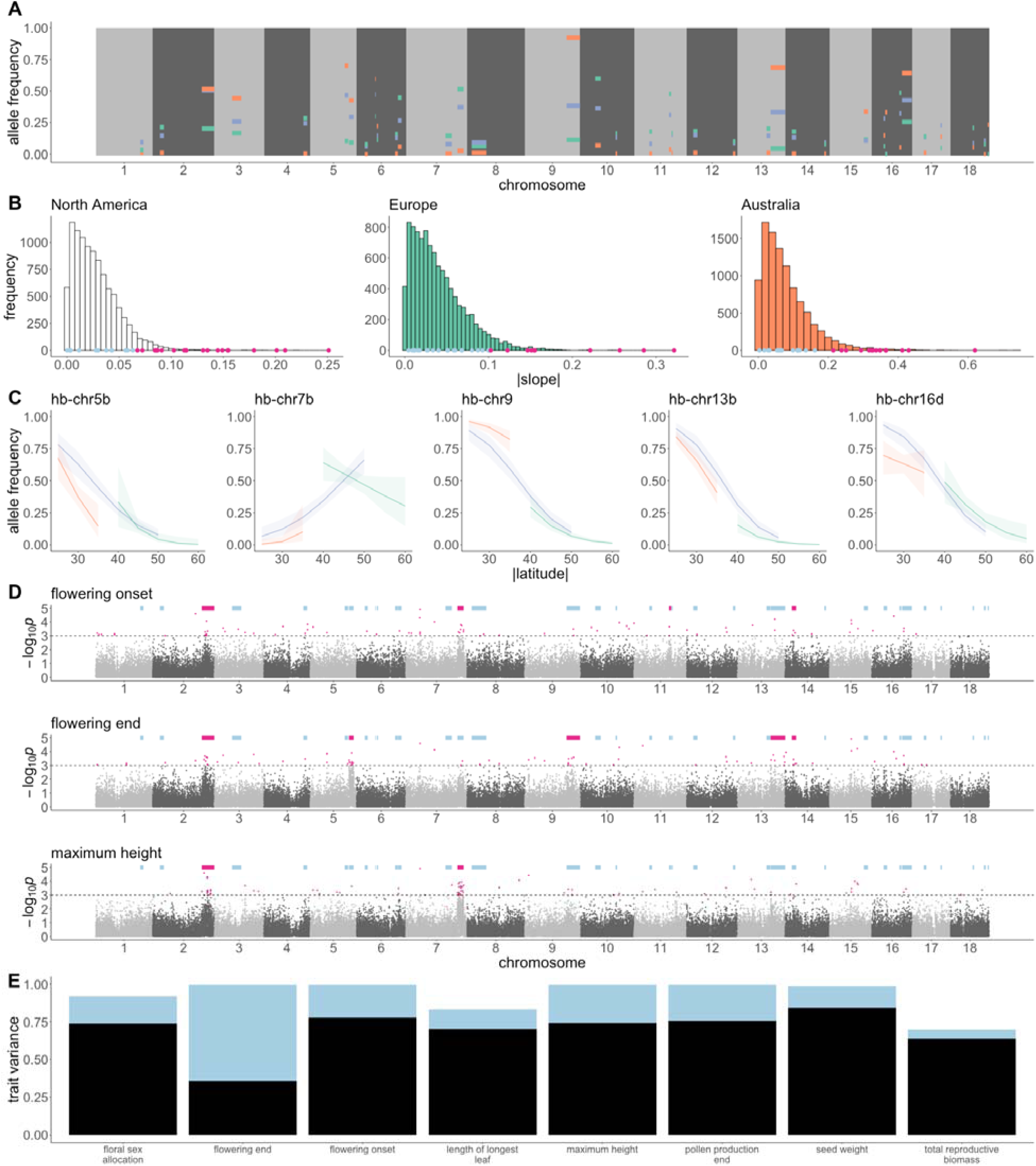
Geographic distributions and phenotypic associations for haploblocks across three ranges (North America: blue; Europe: green; Australia: orange). **A.** Genomic location and frequency of each haploblock in each range. **B.** Distributions of slope estimates of allele frequency as a function of latitude in each range 10,000 random SNPs (histograms) with slope estimates for haploblocks indicated below in blue, or pink for those haploblocks in the 5% tail of the slope distribution. **C.** Logistic regression models with error bands representing 95% CI of least-squares regressions of haploblock frequency against latitude. **D.** Manhattan plots (log-transformed empirical *p*-values for 10 kbp analysis window scores against genomic location) for three phenotypes. Pale blue bars indicate the location of haploblocks. Pink bars denote haploblocks that are enriched for the windows in the 0.1% tail of the *p*-value distribution (indicated by pink points and the dotted line). **E.** Variance explained by different types of genetic variation for the eight traits that showed significant contributions of GWAS outlier-enriched haploblocks (blue) and polygenic background (black).

To assess the involvement of haploblocks in adaptation to local climate, we compared the change in allele frequency in response to latitude for haploblocks and 10,000 randomly sampled SNPs located outside haploblocks and genes. The majority of haploblocks showed evidence of local climate adaptation: of the 37 haploblocks, 30 (81%) were significant (i.e., had slopes in the 5% tail of the slope distribution) in at least one range, and North America, Europe and Australia had 21 (57%), ten (27%) and 18 (47%) significant haploblocks respectively (Fig. 2B; Table S5). Nine haploblocks showed evidence of parallel climate adaptation between North America and at least one invasive range, with slopes significant and in the same direction in each range (Fig. 2C; Table S5). For one haploblock, *hb-chr5b*, parallel patterns were observed between all three ranges (Fig. 2C; Table S5).

To understand the contributions of haploblocks to locally adaptive traits such as size and the timing of flowering, we performed genome-wide association studies using 226 samples from across the three ranges with common-garden phenotypes previously measured by van Boheemen, Atwater and Hodgins (van Boheemen et al., 2019). Ten haploblocks harbored an enrichment of windows (hypergeometric *p*-value<0.05 Bonferroni-corrected for multiple tests across haploblocks) that were associated (0.1% tail of the genome-wide distribution) with at least one of the 13 phenotypes (Table S6). Several haploblocks were associated with multiple traits (e.g., *hb-chr2*; Fig. 2D), and flowering-time phenotypes showed associations with the greatest number of haploblocks (Fig. 2D). All haploblocks associated with flowering-time phenotypes contained *Arabidopsis thaliana* flowering-time gene orthologs (Bouché et al., 2016) (Table S7), but this only constituted a significant enrichment for the 16 orthologs in *hb-chr5b* (Fisher’s exact test *p*-value=0.001; Table S7) which also colocalises with a large-effect flowering time QTL in *A. artemisiifolia* (Prapas et al., 2022). Correspondingly, genes across all haploblock regions were enriched for gene ontology terms related to phenology (e.g., floral organ senescence; regulation of flower development) as well as defense (e.g., regulation of innate immune response; regulation of defense response; Table S8) which are important traits for local adaptation. To understand the specific contribution of haploblocks to these traits, we partitioned for each trait the variance explained by associated haploblocks from the rest of the genomic background represented by a genetic relatedness matrix. Models for ten traits showed support for background additive genetic variance. For eight of these ten traits, a significant proportion of the variance was attributable to genetic effects located in haploblocks, which explained between 14.4% and 63.6% of trait variation (Table S9; Fig. 2E).

### Genomic signatures of selection reveal the extent of parallel evolution

Matching climate associations between native and invasive ranges suggest rapid parallel adaptation has occurred in the invasive ranges to re-establish patterns of local adaptation to climate present in the native range. To address this quantitatively, we first looked for extreme allele frequency divergence between populations within each range as a measure of local adaptation (XtX; Gautier, 2015). We also measured the rank correlation □ (Kendall, 1938) between allele frequency in each range and four independent bioclimatic variables (BIO1: annual mean temperature; BIO2: mean diurnal range; BIO12: annual precipitation; BIO15: precipitation seasonality). For each variable in each range, the 5% tail of the XtX outlier window distribution was significantly enriched for windows in the 5% tail of correlations with each climate variable (*p*<=2.5×10^-23^; Table S10). This suggests that local climate adaptation is a strong driver of allele frequency divergence in each range. Overall, 77.5%, 28.4% and 43.8% of North American, European and Australian XtX outlier windows respectively were also correlated with at least one environmental variable, which is consistent with adaptive differentiation in response to climate. These “XtX-EAA” climate adaptation candidate windows showed significant parallelism between ranges in each pairwise comparison (*p*<=1.37×10^-66^; Fig. 3A; Table S10) and were enriched in 19, six and four haploblock regions in North America, Europe and Australia respectively (Fig. S7; Table S11).

**Figure 3.**
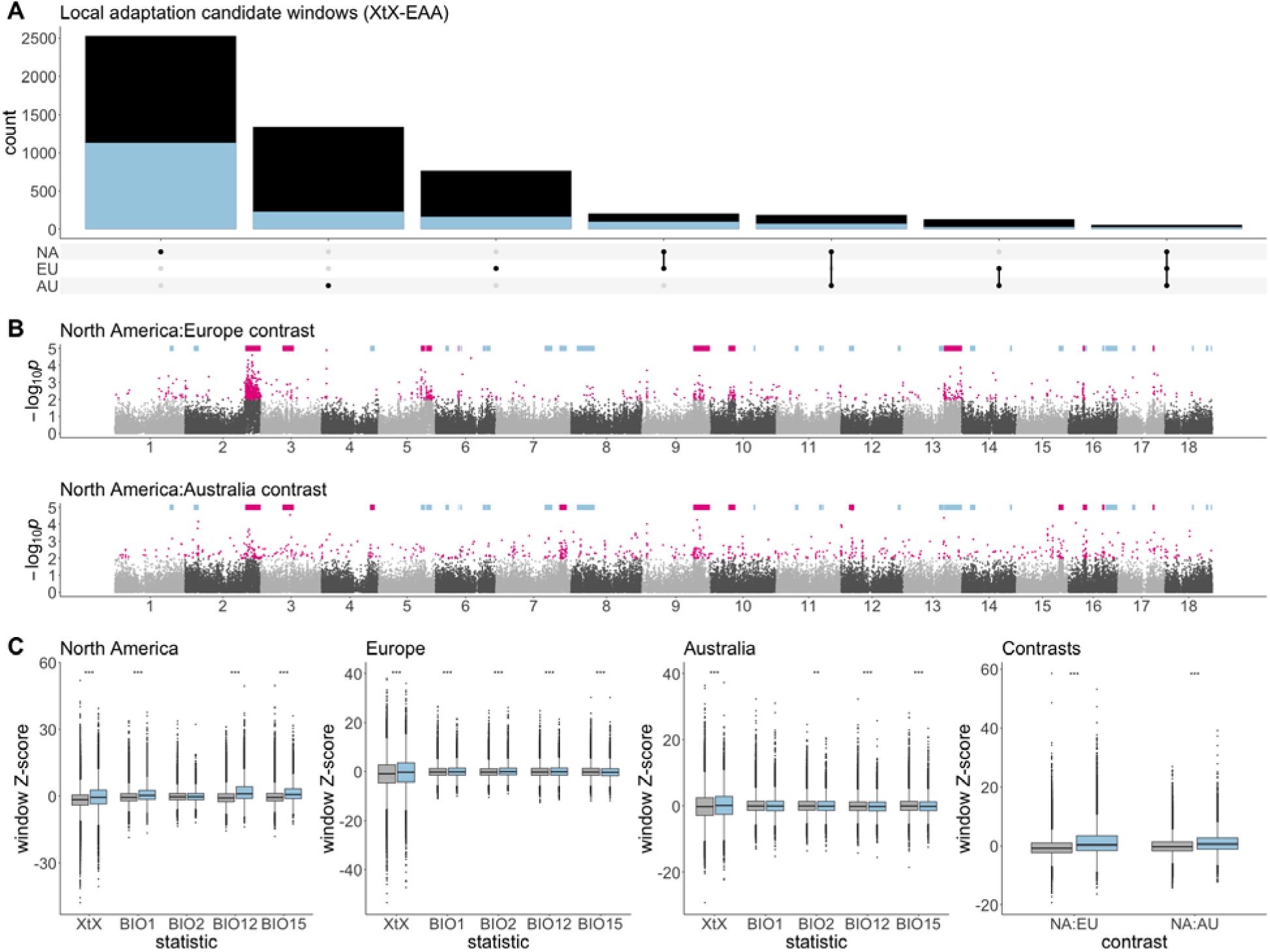
Genomic signatures of parallel rapid adaptation. **A.** Upset plot showing counts of and overlaps between XtX-EAA local adaptation candidate windows in each range (black columns), all of which are enriched for the presence of windows within haploblock regions (blue columns). **B.** Manhattan plots (log-transformed empirical *p*-values for 10 kbp analysis window scores against genomic location) for contrast scans between North America and each invasive range. Pale blue bars indicate the location of haploblocks. Pink bars denote haploblocks that are enriched for the windows in the 1% tail of the *p*-value distribution (indicated by pink points). **C.** *Z*-score distributions of 10 kbp windows for genome scans in haploblock (blue) and non-haploblock (grey) regions of the genome. Asterisks represent *p*-values from Mann Whitney U tests comparing means between haploblock and non-haploblock regions (*: *p*<0.05; **: *p*<0.01; ***: *p*<0.001).

Dramatic shifts in allele frequencies between invasive and native ranges are strong indications of rapid adaptation following introduction. To identify regions of the genome showing such signatures of selection, we computed the BayPass contrast statistic (Olazcuaga et al., 2020) to compare allele frequencies across each invasive range with their likely source in the native North American range separately. We supplemented contrast analyses with selective sweep scans in individual ranges using Fay and Wu’s *H* (Fay and Wu, 2000). Genomic windows showing evidence of native-invasive divergence (contrast outlier windows) showed far more overlap with invasive-range sweep outlier windows (extreme negative *H* values) than those from North America, consistent with rapid adaptation occurring in each introduced range during invasion (3 vs. 96 and 4 vs. 43 for North America:Europe and North America:Australia contrasts respectively; Fig. S8). Among the 1% tail of native-invasive divergence there was significant parallelism between ranges (*p* = 1.44×10^-76^; Table S10). Ten and twelve haploblocks were enriched for contrast outliers in Europe and Australia contrasts respectively, with six (*hb-chr2b*, *hb-chr3*, *hb-chr9*, *hb-chr10a*, *hb-chr16a* and *hb-chr17b*) showing enrichment in both ranges (Fig. 3B; Table S11).

To determine whether regions of the genome harboring large structural variants are more likely to contain candidates for adaptation, we asked whether genomic windows in haploblock regions contained significantly higher values on average (i.e., more evidence for a role in adaptation) than windows in non-haploblock regions of the genome. For all within-range XtX, most correlations with environmental variables, and both native:invasive range contrasts, genome-wide signatures for adaptation were significantly higher in haploblock regions (Fig. 3C; Table S12).

## Discussion

Leveraging whole-genome resequencing of *A. artemisiifolia* samples spanning the native North American range and two introduced ranges in Europe and Australia, we reconstructed the demographic history of these two invasions. Europe was colonized through multiple introductions from across North America, excluding populations from Florida. Contrastingly, most Australian populations appear to be sourced from different parts of North America, particularly Florida. It is often observed that a species’ native and invasive populations occupy similar climate niches (Liu et al., 2020), and the climatic conditions in native environments of *A. artemisiifolia* broadly correspond to those in the invasive ones that they seeded (Fig. S5). Climate matching like this suggests that preadaptation has played a role in determining the success of this invader, and is consistent with predictions from genetically informed species distribution modelling in this species (Putra et al., 2023). However, preadaptation does not preclude the occurrence of post-introduction adaptation (Sherpa and Després, 2021). The results of this study provide compelling empirical evidence of repeated rapid adaptation in *A. artemisiifolia* invasions facilitated by large-effect structural variants introduced from the native range.

Substantial differences exist between the European and Australian invasions of *A. artemisiifolia.* European ragweed occupies diverse climatic conditions and has been present on the continent for over 200 years (Chauvel et al., 2006), while Australian ragweed occupies a more restricted range of climates (Putra et al., 2023; van Boheemen et al., 2019) and was first recorded in Australia in 1908 (Parsons and Cuthbertson, 2001). The results of our demographic analyses highlight further differences between the invasions of Europe and Australia. Notably, the importance of different sources from within the native range to each invasion (Fig. 1C), and the evidence of a strong bottleneck in the Australian, but not the European invasion (Fig. 1E). European results are consistent with previous analyses which suggest that multiple introductions to Europe have prevented or ameliorated the occurrence of a bottleneck during this invasion (Bieker et al., 2022; van Boheemen et al., 2017). Results of a previous analysis of the Australian invasion (van Boheemen et al., 2017) are consistent with our inference of a bottleneck during introduction. In contrast with our results, van Boheemen et al. (2017) inferred the source of the Australian invasion to be within the European range. However they did not extensively sample Florida, a major contributor to the Australian invasion in our analysis, which underscores the importance of exhaustive sampling of native range in demographic studies of invaders.

Invasion events have long been theorized to be associated with population bottlenecks (Baker and Stebbins, 1965), resulting in decreased effective population sizes and reductions in genetic variation (Nei et al., 1975). Bottlenecks may also affect the abundance of deleterious alleles, which can increase due to genetic drift (Whitlock and Davis, 2011), or be purged due to inbreeding (Marchini et al., 2016). Importantly, the drop in Australian effective population size does not appear to have resulted in the accumulation of deleterious alleles (estimated from the average derived allele frequency across zero-fold degenerate sites). These values are similarly low in Australia and the invasion’s North American sources (Fig. 1F), suggesting that the bottleneck has not been long or severe enough to cause an increase in genetic load, despite causing reductions in genetic diversity (van Boheemen et al., 2017; Fig. 1E).

Invasive species introduced to multiple ranges provide a natural experiment to assess the repeatability of adaptation during range expansion (Bock et al., 2015). Repeated genetic evolution reveals constraints and biases in the genetic routes to adaptive optima (Conte et al., 2012; Yeaman et al., 2018), and hence the predictability of adaptation. Furthermore, repeatability suggests the involvement of genetic architectures featuring genetic variants of large effect (Chevin et al., 2010). Population-genomic analyses of parallel invasions are beginning to provide important empirical support in these areas. Repeatability between invasive ranges has been observed in diverse taxa (Battlay, Hendrickson, et al., 2024; Olazcuaga et al., 2020; van Boheemen and Hodgins, 2020), in some cases involving large effect variants (Battlay, Hendrickson, et al., 2024). Our findings, that the invasion histories of *A. artemisiifolia* in Europe and Australia are distinct, allow us to examine the extent to which demography affects these patterns. We have previously described genomic signatures of local climate adaptation in parallel between the native North American range and the invasive European range (Battlay et al., 2023). Here we demonstrate that similar signatures of repeatability are found in the invasion of Australia, and moreover exist between invasions of Europe and Australia. We also observe genomic signatures of adaptive divergence from the native range in parallel between Europe and Australia, reflecting shared genetic changes across multiple independent invasions.

Across our 443 samples we have identified 37 large haploblocks that cover almost 20% of the genome and show features consistent with chromosomal inversions – 15 of these haploblocks correspond to inversion polymorphisms segregating in our diploid reference genome assembly (Fig. S7). Many haploblocks are associated with traits that are important for local climate adaptation (Fig. 2C), and regions harboring haploblocks are enriched for genomic signatures of local adaptation (Fig. 3C). Inversions are predicted to play an important role in local adaptation by maintaining complementary combinations of alleles in blocks of reduced recombination (Kirkpatrick and Barrett, 2015; Kirkpatrick and Barton, 2006). However, whether and under what conditions inversions would support parallel evolution is less clear (Westram et al., 2022). Nevertheless, haploblocks appear to play a key role in parallel evolution in this species. By leveraging common garden data from samples spanning all three ranges, we observe that haploblocks underpin traits important for adaptation to local climate. In many cases haploblock allele frequency clines present in the native range have rapidly reformed in invasive ranges (Fig. 2B; Fig. 2C). Haploblocks are also disproportionately represented in parallel divergence between native and invasive ranges. Of the 98 outlier windows shared between North America:Europe and North America:Australia contrasts, 70 occur in haploblocks (*p* = 7.79×10^-20^). The haploblocks of *A. artemisiifolia* are predominantly (if not completely) standing variation in the native range, where we observe in our data segregation of 36 of the 37 haploblocks (Fig. 2A). The observed strong signatures of selection on these variants in Australia adds to mounting empirical evidence that bottlenecks do not necessarily limit adaptation or need to be overcome by *de novo* mutations (e.g., Dogantzis et al., 2024).

As the number of biological invasions worldwide continues to grow, understanding the extent, genetic architecture and predictability of range expansion in invasive species becomes ever more salient. Our results showcase the remarkable parallel adaptation that has occurred as demographically distinct introductions of *A. artemisiifolia* have invaded Europe and Australia over the last 200 years. Central to this result are 37 large structural variants, which underscore the role of large-effect standing variation in rapid and repeated adaptation during invasion.

## Methods

### Ambrosia artemisiifolia samples

We generated whole-genome resequencing data for 95 specimens of *A. artemisiifolia* collected from the east coast of Australia in 2014 and described in (van Boheemen et al., 2019). We combined this data with whole-genome resequencing of 348 *A. artemisiifolia* specimens collected from North America (*n*=179) and Europe (*n*=169) between 2007 and 2019 and previously described in (Bieker et al., 2022). See Table S1 for details of each sample.

### Whole-genome resequencing

Illumina libraries were prepared following the approach of (Bieker et al., 2022). Leaf tissue was collected and stored in silica gel desiccants (Chase and Hills, 1991) at room temperature until required for DNA extraction. Approximately 20 to 30 mg of dried leaf tissue from each sample were placed inside a 2.0-ml tube with a 3-mm stainless steel bead and ground with a TissueLyser II (QIAGEN). The DNA was extracted using a modified CTAB protocol (Doyle and Doyle, 1987) adapted for a 96-well plate format (Ivanova et al., 2008) using EconoSpin filter plates, and the DNA was suspended in 60 μl of elution buffer. Extracted DNA was quantified using a Qubit 2.0 fluorometer (Invitrogen, Carlsbad, CA, USA) using dsDNA Quantitation HS (high sensitivity, 0.2 to 120 ng) kit. Extraction blanks were prepared alongside the samples to monitor possible contamination. Extracts were converted into blunt-end Illumina libraries as described above using BEST protocol (Carøe et al., 2018), in which custom blunt-end adapters (Meyer and Kircher, 2010) were ligated to the DNA fragments or single-stranded Illumina libraries using the Santa Cruz Reaction protocol (Kapp et al., 2021). Dual-index libraries were generated using custom index primers during indexing PCR. Indexing PCR was carried out in a 50-μl reaction using 10 μl of library template, 0.2 μM sample-specific forward index primer, 0.2 μM sample-specific reverse index primer, 1× Platinum SuperFi PCR master mix and the rest of the volume filled up with molecular-grade water. The PCR was performed with an initial denaturation of 3 min at 95°C, followed by 12 cycles of a 20-s denaturation at 98°C, 60 s of annealing at 60°C, and 60 s of extension at 72°C, followed by a final extension for 5 min at 72°C. Amplified libraries were purified with SPRI beads (Rohland and Reich, 2012) and eluted in 33 μl of EBT buffer. The samples were sent to Genewiz and sequenced on one lane of Illumina NovaSeq 6000 PE150 which yielded 935 Gbp of data.

### Alignment and variant calling

Each pair of FASTQ files from 95 Australian *A. artemisiifolia* samples were aligned to the primary haplotype of the phased, diploid reference assembly described in (Battlay et al., 2023) using the *Paleomix* pipeline v.1.2.13.4 (Schubert et al., 2014), which incorporates *AdapterRemoval* v.2.3.1 (Schubert et al., 2016), *BWA-MEM* v.0.7.17 (Li, 2013), *Picard* v.2.19.0 *MarkDuplicates* (https://broadinstitute.github.io/picard/) and GATK v.3.7 *IndelRealigner* (Van der Auwera and O’Connor, 2020). Mean depths of alignments ranged from 1.2X to 15.6X with a mean of 5X (Table S1). Variants were called across the resulting 95 BAM files from Australian samples and 348 BAM files previously generated from North American and European samples (Battlay et al., 2023; Bieker et al., 2022). GATK v.3.8 *UnifiedGenotyper* (DePristo et al., 2011) was used to call variants on all contigs greater than 100 kbp in length. GATK v.3.8 *VariantFiltration* (Van der Auwera and O’Connor, 2020) and *VcfTools* v.0.1.16 (Danecek et al., 2011) were used to filter variant calls. SNP and indel calls were separately filtered using GATK hard-filtering recommendations (SNPs: QD□<□2.0, FS□>□60.0, SOR□>□3.0, MQ□<□40.0, ReadPosRankSum□<□−8.0, MQRankSum□<□−12.5; indels: QD□<□2.0, FS□>□200.0, SOR□>□10.0, ReadPosRankSum□<□−20.0, InbreedingCoeff□<−0.8). In addition, SNPs and indels were separately filtered for sites with depth (DP) less than one standard deviation below the mean, and greater than 1.5 standard deviations above the mean. Individual genotypes were set to missing if their depth was less than three, then variants with greater than 20% missing across all samples were removed. Samples with greater than 60% missing variants were removed. For the remaining 397 samples, genotypes were phased and imputed using Beagle v.5.4 (Browning et al., 2018).

### Population structure

To calculate genotype likelihoods we ran *angsd* v0.939 (Korneliussen et al., 2014; -GL 2 - doMajorMinor 1 -doCounts 1 -doGLF 2 -SNP_pval 1e-6 -doMaf 2 -doGeno -1 -doPost 1 -minMapQ 30 -minQ 20 -trim 5 -minMaf 0.05 -geno_minDepth 2 - setMinDepthInd 2 -uniqueOnly 1 -doPlink 2) on alignments of 443 samples, including only sites that could be estimated in >75% of samples (-minInd 333). A list of linkage disequilibrium (LD)-pruned variants was generated in *plink* v1.9 (Chang et al., 2015; -- indep-pairwise 50 5 0.5). Sites were filtered to include only those that were LD-pruned and also outside annotated genes and haploblocks, and then randomly downsampled to 100,000. A covariance matrix was generated from these filtered sites using *pcangsd* v.1.2 (Meisner and Albrechtsen, 2018) and used for principal component analysis (PCA) in R v.4.3.1. Admixture analysis was performed using *NGSadmix* v.0.939 (Skotte et al., 2013) and the same genotype likelihoods as the PCA. We ran analyses for two to six ancestral populations (*K*). Ten independent runs with different seeds were performed for each *K* value. The run with the highest likelihood for each *K* was used for plotting (Fig. S2).

### Demographic modeling

We implemented an approximate Bayesian computation (ABC) random forest (RF) statistical framework using *DIYABC-RF* (Collin et al., 2021) to model the formation of genetic structure in the native range, infer the introduction source of the invasive ranges, and to estimate demographic parameters of *A. artemisiifolia*. For this ABC-RF analysis we divided North America into the four clusters previously defined (Bieker et al., 2022) based on *ADMIXTURE* results and geography. We used all samples for which SNPs were called, excepting the southernmost Australian population, which appeared separate to the main genetic cluster (AU01; Fig. S1). SNPs were LD-pruned with *plink* 2.0 alpha (Chang et al., 2015; `indep-pairwise 50 5 0.2`) and filtered for MAF>0.05. We built on the ABC-RF analysis of van Boheemen et al. (2017), modeling both the development of genetic structure in the native range in addition to incorporating independent introduction and bridgehead introduction scenarios for each invasive range (Fig. S3; Fig. S4). For the introduction scenarios, we used the priors defined in (van Boheemen et al., 2017; (Table S13), to which readers are directed for detailed description.

To model the formation of genetic structure in the native range of *A. artemisiifolia*, we simulated datasets from nine branching pattern topologies, each with 3 temporal parameter configurations, for all permutations of the four sampled native range populations (Fig. S3). To manage the number of native range scenarios to be compared with ABC-RF we opted for a sequential approach to model selection. This approach first compared subsets of scenarios - defined by topology - before comparing the best-supported scenarios between subsets to determine the overall best scenario (Fig. S3; Table S2; Byrne et al., 2022). In a stepwise manner we conducted independent analyses of the introduction of *A. artemisiifolia* to Europe and Australia, incorporating the results of the preceding native range analysis (Fig. S4; Table S3; Fontaine et al., 2021). In the final analysis we compared the best-supported scenarios of independent introduction with bridgehead introductions from Europe to Australia (Table S14). Further details of the implementation of this method are provided in the supplementary materials.

### Invasion source prediction

To predict source locations of invasive *A. artemisiifolia* populations, we used *Locator* (Battey et al., 2020) and SNP data from the 397 samples for which SNPs were called (Table S1). All SNPs with MAF>0.05 were included in the analysis. Models were trained using sampling location (latitude; longitude) and genotype data for all samples across the putative source range and then the locations of invasive-range individuals were predicted by the model. For Europe we used North America as the source range. For Australian samples we initially included North America and Europe as sources, but as all Australian source locations in this pilot run were predicted to be much closer to North America than Europe, we subsequently used only the North American range as a source for Australian samples. We ran Locator in 10-Mbp non-overlapping windows across the genome, and averaged each prediction location across these 113 genomic windows.

### Effective population size

We used *GONE* (Novo et al., 2023) to estimate effective population size in each range over time. In North America and Europe we used all samples for which SNPs were called (*n*=155 for each range). In Australia we only included the 78 samples that were in the main genetic cluster (see Fig. 1B; Table S1) to avoid the confounding effects of population structure on estimates of effective population size (Novo et al., 2023). In each range and for each North American cluster, *GONE* was run using 100,000 SNPs sampled from outside haploblock regions, with the recombination rate parameter was set to 1.428 cM/Mbp based on the genetic map described by Prapas et al. (2022).

### Estimation of genetic load

To identify zero-fold and four-fold degenerate sites throughout the genome, we developed a Python script to systematically examine all possible codons (based on the reference annotation; Battlay et al., 2023) for substitutions that always (zero-fold degenerate) or never (four-fold degenerate) result in amino acid substitutions. We analyzed each invasive range and each North American genetic cluster separately. Within each range or cluster we randomly downsampled the number of individuals to equal the number of individuals in the group with the smallest sample size (North America [south]; *n*=22) and used *angsd* v.0.939 (Korneliussen et al., 2014) to calculate the frequency of derived sites in each group (`-GL 1 -doMajorMinor 5 -doMaf 2 -minQ 20 -minMapQ 30 -minInd 4 -trim 5`), using the consensus of resequencing data from two outgroup species (*Ambrosia chamissonis* and *Ambrosia carduacea*; Bieker et al., 2022) mapped to the reference to determine the ancestral state at each site (`-anc`). We calculated the mean frequencies of genome-wide derived zero-fold and four-fold degenerate mutations for each group and performed bootstrapping by drawing 20% of sites for each group to recalculate means, which was repeated 100 times.

### Haploblock identification

We identified haploblocks (indicative of large structural variants) using local principal component analysis, modifying the method described by Li and Ralph (2019) to utilize covariance matrices from *pcangsd* v.1.2 (Meisner and Albrechtsen, 2018), which were calculated in 100-kbp windows from beagle files generated in *angsd* v.0.939 (Korneliussen et al., 2014; `-GL 2 - doMajorMinor 1 -doCounts 1 -doGLF 2 -SNP_pval 1e-6 -doMaf 2 -doGeno -1 -doPost 1 -minMapQ 30 -minQ 20 -trim 5 -minMaf 0.05 -geno_minDepth 2 - setMinDepthInd 2 -uniqueOnly 1 -doPlink 2`) on alignments of 443 samples, including only sites that could be estimated in >75% of samples (`-minInd 333`). Local population structure along each chromosome was analyzed on five MDS axes and outliers were identified from the 5% corners of each pair of MDS axes. Candidate haploblocks were identified by manual examination of MDS plots for each chromosome. Across each candidate region local PCAs were performed with *angsd* and *pcangsd* using the same parameters used for the 100kbp windows. Heterozygosity was also calculated for each sample in each candidate region using *angsd* (`-dosaf 1 -minMapQ 30 -minQ 20 -trim 5 -GL 2`) and *realSFS* v.0.939 (Nielsen et al., 2012; `-fold 1`). Haploblocks were identified by the presence of three clusters of samples along a single principal component axis, indicative of two homozygous and one heterozygous inversion genotype, as well as by the presence of elevated heterozygosity in the region for samples genotyped as heterozygotes. We defined elevated heterozygosity as when the standard error of the mean (SEM) for each homozygote class did not overlap the SEM for the heterozygote class. We validated haploblock genotypes by performing LD scans with *ngsLD* v.1.2.0 (Fox et al., 2019; `--min_maf 0.05 --max_kb_dist 0`) on 5,000 randomly sampled sites from each chromosome containing a haploblock. For each haploblock, LD scans were run on all samples, as well as a set of samples homozygous for the more common haploblock allele. To identify inversions segregating in our diploid reference that corresponded to population-genomic haploblock signatures, we aligned the two haplotypes of our reference (Battlay et al., 2023) using *minimap* v.2.1.8 (Li, 2018; `-t 12 -P -k19 -w19 -m200`). After filtering out alignments that were shorter than 10 kbp or that contained fewer than 5,000 matches, we visualized *minimap* alignments using the R package *pafr* and manually compared them to MDS manhattan plots of local PCA results (Fig. S6).

### Haploblock-latitude slopes

To identify haploblocks exhibiting clinal patterns, we used generalized linear models to assess how the allele frequency of each haploblock (and 10,000 SNPs sampled from outside haploblocks and genes) changed with latitude in each range separately. We designated significant slopes as those slopes that fell into the 5% tail of the SNP distribution. We also ran generalized linear models across all three ranges to assess how haploblock allele frequency changed with absolute latitude, using range (North America, Europe or Australia) and latitude as well as their interaction as fixed effects. Non-significant interactions were removed. Model estimates along with the 95% CI ribbons were plotted with the R package *emmeans* (Lenth et al., 2019).

### Genome-wide association

Of the 443 samples in this study, 226 had been measured in a common garden for multiple ecologically important phenotypes by van Boheemen et al. (2019). We performed genome-wide association studies (GWAS) using 13 phenotypes: *floral sex allocation, flowering end, flowering onset, length of longest leaf, length of longest raceme, maximum height, pollen production end, number of racemes, root/shoot ratio, seed weight, shoot biomass, total biomass,* and *total reproductive biomass*. Three outlier sample measurements (> three standard deviations from the mean) were removed from the *length of longest leaf* phenotype. *Floral sex allocation* and *number of racemes* were respectively log_10_- and square root-transformed.

Beagle files for GWAS were generated in *angsd* v0.939 (Korneliussen et al., 2014) using the same parameters as the population structure analysis (`-GL 2 -doMajorMinor 1 - doCounts 1 -doGLF 2 -SNP_pval 1e-6 -doMaf 2 -doGeno -1 -doPost 1 - minMapQ 30 -minQ 20 -trim 5 -minMaf 0.05 -geno_minDepth 2 - setMinDepthInd 2 -uniqueOnly 1 -doPlink 2`) and using only sites that could be estimated in >75% of samples (`-minInd`). GWAS were performed in *angsd* v.0.939 (Korneliussen et al., 2014; `-yQuant -cov -Pvalue 1 -doAsso 4`). To account for population structure we used the first two principal components of a covariance matrix generated using *pcangsd* v.1.2 (Meisner and Albrechtsen, 2018) using the same parameters as the population structure analysis: a sample of 100,000 LD-pruned variants (*plink* v1.9; Chang et al., 2015; `--indep-pairwise 50 5 0.5`) from outside annotated genes and haploblocks.

GWAS results were analyzed in 10-kbp non-overlapping windows across the genome using *The Weighted-Z Analysis* (*The WZA*; Booker et al., 2023).

### Gene ontology and flowering time gene enrichment

Gene ontology (GO) enrichment was assessed using functional annotations described in Battlay et al. (2023). To identify GO terms enriched among candidate lists, the R package *topGO* (Alexa and Rahnenführer, 2009) was used with Fisher’s exact test, the ‘weight01’ algorithm, and a *p*-value<0.05 to assess significance. In addition, we assessed enrichments of the 513 *A. artemisiifolia* genes that are orthologs of *A. thaliana* FLOR-ID flowering time pathway genes (Bouché et al., 2016) using Fisher’s exact test and a *p*<0.05 threshold.

### Trait variance partitioning

To partition trait variation into contributions from haploblocks vs the remaining polygenic background (Koch et al., 2022), we built a genomic relationship matrix (GRM) from 10,000 LD-thinned SNPs outside haploblocks using the R package *AGHMatrix* (Amadeu et al., 2023). We used SNPs from the 207 samples with both SNP and phenotype data available (Table S1), and made the matrix positive-definite by adding a small positive value (1×10^-15^) to one eigenvalue smaller than it using the R package *mbend* (Nilforooshan, 2020).

Next, we fitted a model to each trait in a Bayesian framework as implemented in the R package *MCMCglmm* (Hadfield, 2010). Each model included genotypes between one and five trait-associated haploblocks as fixed effects, and the genomic relationship matrix as a random effect to model the effect of additive genetic variation located outside haploblocks. We set a parameter-expanded prior on random effects (*V*=1, Ν=2, □ =0, □ =1000) and an Inverse-Wishart prior (*V*=1, Ν=0.002) on residuals. 7,010,000 Markov chain Monte Carlo iterations were run with a thinning period of 7,000 iterations and a burn-in period of 10,000 iterations. Model parameters were estimated from their posterior distribution using 1,000 thinned iterations. Autocorrelations between MCMC samples were below the recommended level of 0.1, yielding effective sample sizes >753 for all estimates. We inspected plots of traces and posterior distributions to ensure that models converged. Other priors were explored and gave similar results to those presented here.

Last, we calculated the total variation of each trait as the sum of the variation explained by haploblocks (calculated following de Villemereuil et al. [2018]), the additive genetic variation outside haploblocks, and the residual variation. We then calculated the proportion of total trait variation explained by each component, along with its 95% credible interval.

### Local adaptation scans

For each sampling location we extracted 19 bioclimatic variables from the WorldClim database (Fick and Hijmans, 2017) using the R package *raster* (Hijmans et al., 2015), and selected four bioclimatic variables (BIO1: annual mean temperature; BIO2: mean diurnal range; BIO12: annual precipitation; BIO15: precipitation seasonality) that were minimally correlated (*r*<0.7) in all three ranges for further analysis. We first generated beagle files in *angsd* v.0.939 (Korneliussen et al., 2014; `-GL 2 -doMajorMinor 1 -doCounts 1 -doGLF 2 - SNP_pval 1e-6 -doMaf 2 -doGeno -1 -doPost 1 -minMapQ 30 -minQ 20 -trim 5 -minMaf 0.05 -geno_minDepth 2 -setMinDepthInd 2 -uniqueOnly 1`) in each range separately using all samples from populations with *n*>2, and with the `-minInd` flag set to 75% of the number of samples per range. The resulting sites with rangewide allele frequencies >0.05 were then measured in each population separately in subsequent *angsd* analyses (`- doMaf 4 -minInd 2`) with no minor allele frequency filter. The *BayPass* v.2.2 core model (Gautier, 2015; with an Ω covariance matrix computed from 10,000 randomly-sampled sites that were located outside annotated genes and haploblocks) was run using population allele frequencies in each range separately. Correlations between population allele frequencies and the four independent bioclimatic variables were also measured in each range separately using □ (Kendall, 1938). The results of *BayPass* XtX and T correlations were further analyzed for enrichment in 10kbp non-overlapping windows across the genome using *The WZA* (Booker et al., 2023).

### Contrast scans

To identify regions on the genome that have been the target of divergent selection during range expansion, we used the BayPass contrast statistic (Olazcuaga et al., 2020) to compare population allele frequencies across each invasive range with source populations in the native North American range identified by our demographic analyses (Fig. 1; Table S1). We first generated beagle files in *angsd* v.0.939 (Korneliussen et al., 2014; `-GL 2 -doMajorMinor 1 -doCounts 1 -doGLF 2 -SNP_pval 1e-6 -doMaf 2 -doGeno -1 -doPost 1 - minMapQ 30 -minQ 20 -trim 5 -minMaf 0.05 -geno_minDepth 2 - setMinDepthInd 2 -uniqueOnly 1`) in each combination of North American and European or Australian ranges. All samples from populations with *n*>2 were included with the exception of Australia, where we excluded samples presumed to be part of a second, very recent introduction (Fig. S1). For each analysis the `-minInd` flag was set to 75% of the number of samples per range pair. The resulting sites that had frequencies of >0.05 across each range pair were then measured in each population separately with no minor allele frequency filter (`- doMaf 4 -minInd 2`). The *BayPass* v.2.2 contrast model (Olazcuaga et al., 2020) was run comparing population allele frequencies in North America with each of Europe and Australia separately using an Ω covariance matrix computed from 10,000 randomly-sampled sites that were located outside annotated genes and haploblocks.

Contrast scans were conducted in BayPass for each invasive range separately using an omega matrix calculated from 10,000 putatively neutral sites to account for population structure. We assessed the enrichment of extreme contrast values in non-overlapping 10-kbp windows using *The WZA* (Booker et al., 2023). To understand the source of divergence observed in contrast scans, we also scanned each range for evidence of selective sweeps. We calculated *H* (Fay and Wu, 2000) with *angsd* v.0.939 (`-doSaf 1 -doMajorMinor 4 -GL 2 -baq 2 - minMapQ 30 -minQ 20`) using only sites that could be estimated in >75% of samples (`- minInd`), and the consensus of resequencing data from two outgroup species (*Ambrosia chamissonis* and *Ambrosia carduacea*; Bieker et al., 2022) mapped to the reference to determine the ancestral state at each site (`-anc`). *H* was calculated in 10-kbp non-overlapping windows across the genome.

### Data availability

The diploid reference genome assembly used in this study is available from NCBI under BioProject IDs PRJNA929657 and PRJNA929658. Individual sample resequencing data are available from ENA under BioProject IDs PRJEB48563, PRJNA339123 and PRJEB34825, and from SRA under BioProject ID PRJNA1139307. Phenotype data is available from github.com/lotteanna. Code is available from github.com/pbattlay.

## Supplementary Methods

### Demographic modeling

We implemented an approximate Bayesian computation (ABC) random forest (RF) statistical framework using *DIYABC-RF* (Collin et al., 2021) to model the formation of genetic structure in the native native, infer the introduction source of the invasive ranges, and to estimate demographic parameters of *A. artemisiifolia*. In this framework, we simulated genetic data under various demographic scenarios based on user-defined priors (Table S13). We then compared summary statistics from these simulated data with those from the observed data (Beaumont, 2010; Beaumont et al., 2002).

RF, a machine-learning algorithm, constructs decision trees from bootstrapped samples to perform classification tasks. These trees use summary statistics as predictor variables (Breiman, 2001), allowing RF to identify the scenario that most closely aligns with the observed data. *DIYABC-RF* uses all available summary statistics (Table S15), supplemented with axes from linear discriminant analysis (LDA; Pudlo et al., 2016). The RF method reserves a portion of the simulated data as ‘out-of-bag’. This data is used to estimate the error rate, providing a measure of accuracy comparable to using a test set of the same size as the training set (Pudlo et al., 2016).

We used the six genetic units. Four of these units represent different regions in the native North American range and were previously defined by Bieker et al. (2022) based on ADMIXTURE results and geography: (1) south (NA-S); (2) mid east (NA-ME); (3) east (NA-E); (4) western (NA-W). The remaining two genetic units were from the invasive ranges: (5) Europe (EU) and (6) Australia (AU). To account for potential unsampled sources of introductions, we included unsampled or ‘ghost’ genetic units (Slatkin, 2005). These ‘ghost’ genetic units were used to define scenarios modeling multiple introductions, including repeated introductions from the same native source, leading to subsequent admixture in the introduced range.

We used all samples for which SNPs were called (*n*=391), except the southernmost Australian population, which appeared separate to the main genetic cluster (AU01; Fig. S1). All SNPs with MAF>0.05 were included in the analysis. Using plink 2.0 alpha (Chang et al., 2015), all SNPs were filtered to include only those that were LD pruned (`--indep-pairwise 50 5 0.2`) and polymorphic (`--geno 1`). Resulting filtered VCF files were converted to ‘.snp’ format suitable for *DIYABC-RF* analysis using a custom script.

We extended the ABC-RF analysis by van Boheemen et al. (2017), incorporating independent introduction and bridgehead introduction scenarios for each invasive range. These scenarios included founding, secondary, and bottlenecked introductions. We modeled admixture between genetic units both before and after introduction. For the introduction scenarios, we used the priors defined in van Boheemen et al. (2017; Table S13).

We first modeled the divergence of the genetic clusters in the native range. Due to uncertainty around the values of historical priors, we adjusted the lower and upper prior limits based on the prior distributions of the preliminary simulated datasets to ensure a reasonable compatibility between the observed and simulated datasets (Collin et al., 2021). To achieve this we used DIYABC-RF’s visual and numerical outputs, including the projection of the datasets on the first linear discriminant analysis (LDA) axes and the proportion of simulated data with summary statistics values below those of the observed dataset (Collin et al., 2021). This allowed us to set the prior distribution as wide as possible, accommodating our uncertainty, while remaining within biological reason (Bertorelle et al., 2010).

To model the formation of genetic structure in the native range of *A. artemisiifolia*, we simulated datasets from nine branching pattern topologies (Fig. S3). Each topology contains three temporal parameters (*t_anc_*, *t_1_* or *t_4_*, *t_2_*or *t_5_*), within the prior interval of which branching events could occur by divergence or admixture (Table S2). These parameters were configured in three ways: all branching occurring early (*t_1_* and *t_2_*), a combination of early and recent branching (*t_1_* and *t_4_*), or all branching events occurring recently (*t_4_* and *t_5_*). For each topology we considered all configurations of temporal parameters, and all permutations of the four sampled native range populations. This resulted in the generation of 648 unique scenarios (9 topologies × 3 configurations × 4! permutations).

To manage the number of native range scenarios to be compared with ABC-RF we opted for a sequential approach to model selection (Table S2; Table S3; Table S14; Byrne et al., 2022). Initial model choice analyses were conducted on scenarios within their respective topologies (Table S2, analyses 1-9). Scenarios that achieved an above-average number of classification votes (i.e., no. RF trees / no. scenarios) within their topology were then competed across topologies (Table S2, analysis 10). This ensured that well-supported scenarios were directly compared. To determine model choice consistency, we replicated these analyses with independent simulated datasets, using a larger subset of available SNPs and a higher number or RF classification trees (Chapuis et al., 2020).

We conducted independent, stepwise analyses of the introduction of *A. artemisiifolia* to Europe and Australia, incorporating the results of the preceding native range analysis (Fig. S4; Table S3). The native range scenario with the highest posterior probability from the across-topology native range analysis was used as the ancestral scenario for the primary and/or secondary introductions (Fontaine et al., 2021).

We compared scenarios of independent and bridgehead introductions by generating 64 combination scenarios from independent introduction scenarios that received above-average classification votes (Table S3). These were compared against 8 scenarios consisting of above-average independent European introduction scenarios given a bridgehead introduction to Australia (Table S14). Model choice analysis was conducted with an increasing training set (starting at 5,000 per scenario) and trees (starting at 30,000) until model choice converged for eight consecutive runs. The proportion of votes, global prior error rate and posterior probability values are averaged over the replicates.

To assess if scenario choice was sensitive to the number of bridgehead vs. non-bridgehead scenarios, we conducted model choice analysis on a subset of scenarios. In this analysis, the scenario for the non-bridgehead introduction to Australia was based on the scenario with the highest posterior probability from the independent Australian introduction analysis. This scenario was combined with the 8 above-average European introduction scenarios and then competed against an equal number of bridgehead introduction scenarios.

## Supplementary Tables

https://www.dropbox.com/scl/fi/7lne5mlq1ms8q01rz94v8/RagweedAus_STables_bioRxiv.xlsx?key=ibqk6meqf0uvjlamemriwl1wd&dl=1

## Author contributions

**Paul Battlay:** Conceptualization, Data Curation, Formal Analysis, Methodology, Software, Visualization, Writing – Original Draft Preparation, Writing – Review & Editing

**Samuel Craig:** Formal Analysis, Software, Visualization, Writing – Original Draft Preparation

**Andhika R. Putra:** Formal Analysis, Investigation, Software, Writing – Original Draft Preparation

**Keyne Monro:** Formal Analysis, Investigation, Software, Writing – Original Draft Preparation

**Nissanka P De Silva:** Formal Analysis, Investigation, Software, Writing – Original Draft Preparation

**Jonathan Wilson:** Formal Analysis, Software

**Vanessa C. Bieker:** Investigation

**Saila Kabir:** Investigation

**Nawar Shamaya:** Investigation

**Lotte van Boheemen:** Investigation

**Loren H. Rieseberg:** Resources

**John R. Stinchcombe:** Conceptualization, Funding Acquisition, Resources

**Alexandre Fournier-Level:** Conceptualization, Funding Acquisition, Resources, Supervision

**Michael D. Martin:** Conceptualization, Funding Acquisition, Resources, Supervision

**Kathryn A. Hodgins:** Conceptualization, Data Curation, Funding Acquisition, Methodology, Project Administration, Resources, Supervision, Writing – Review & Editing

---

**Figure S1.**
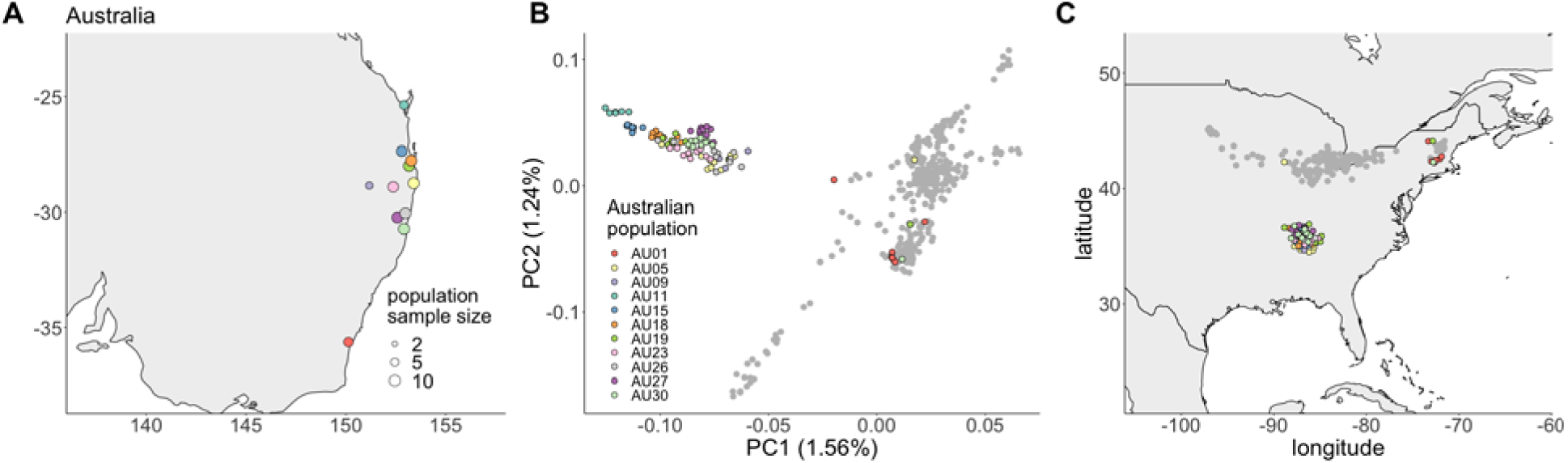
**A**. Geographic location of each Australian population. **B.** The first two principal components of putatively neutral genetic variation for each sample (as in Fig. 1B), with individual Australian samples coloured by population and North American and European samples in grey. **C**. Deep neural network-predicted source locations in North America for invasive-range samples (as in Fig. 1D) with European predictions in grey and Australian predictions coloured by population with colours corresponding to (A).

**Figure S2.**
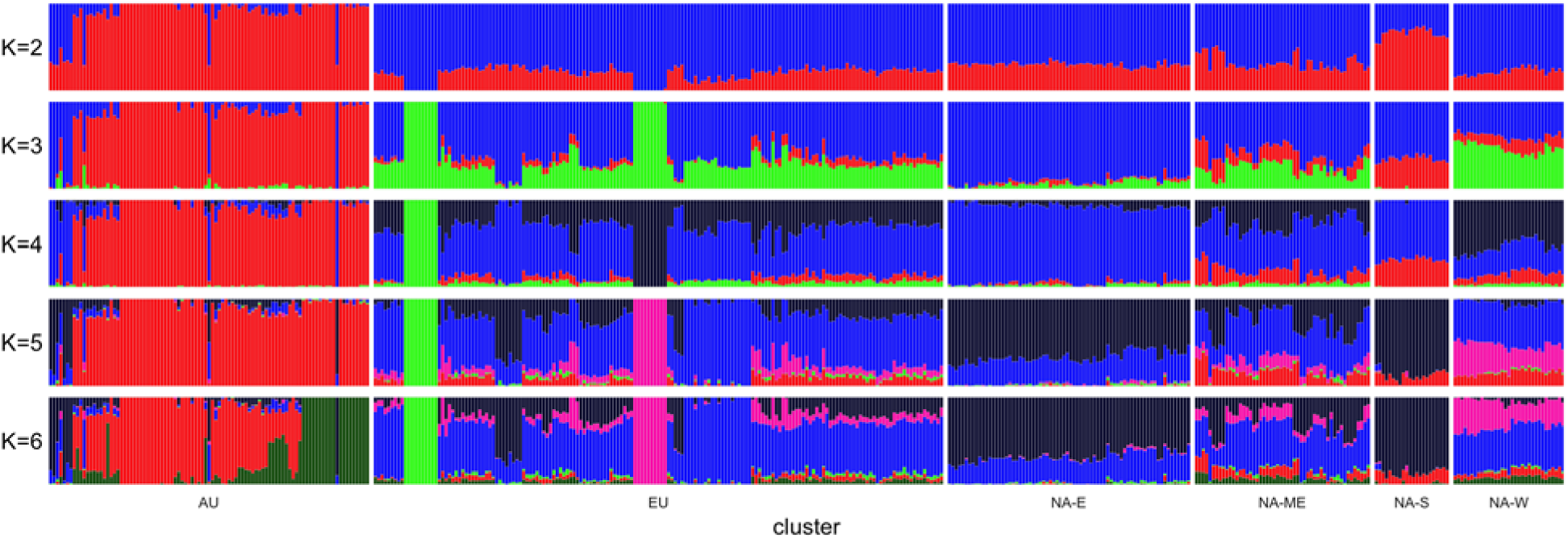
Admixture proportions for 443 *Ambrosia artemisiifolia* samples (vertical bars) for between two and six theoretical ancestral populations (*K*). North American and European samples are arranged by genetic-spatial clusters described in Bieker et al. (2022). Within clusters samples are grouped by sampling location. AU: Australia; EU: Europe; NA-E: North America (east); NA-ME: North America (mid east); NA-S: North America (south); NA-W: North America (west).

**Figure S3.**
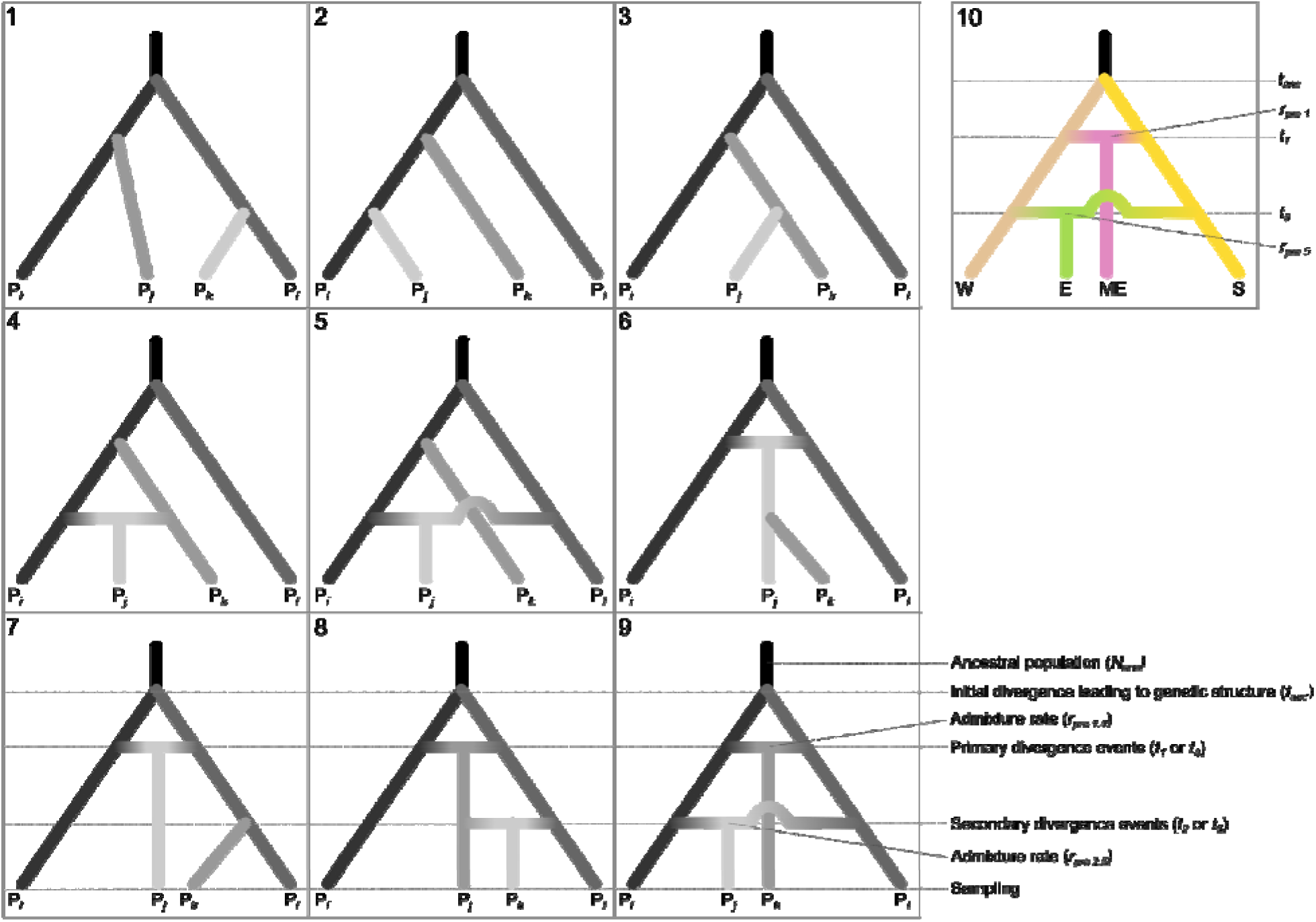
Graphical illustration of the development of genetic structure of *Ambrosia artemisiifolia* within its native range, tested using ABC-RF. For analyses 1-9, P*_i_*, P*_j_*, P*_k_*, P*_l_*represent the four sampled native range genetic units (W, ME, E, S). Scenarios within analyses 1-9 were generated by permuting the four genetic units at the branch tips for each of three divergence event timing configurations: both occurring early (*t_1_* and *t_2_*); a combination of early and recent divergence events (*t_1_* and *t_5_*), or; both occurring recently (*t_4_* and *t_5_*). Effective population sizes, *N_i_*, are represented in grayscale, with the ‘ancestral population’, *N_anc_*, labeled and shown in black. For scenarios where divergence is due to one or more admixture events, two independent pre-introduction admixture rates (*rpre_t1_*_,*t4*_, *rpre_t2_*_,*t5*_) were modeled. The best-supported scenario from analysis 10–which compared well-supported scenarios from analyses 1-9– is shown in color. Time is not to scale. For detailed descriptions of parameters and prior intervals, refer to Table S13.

**Figure S4.**
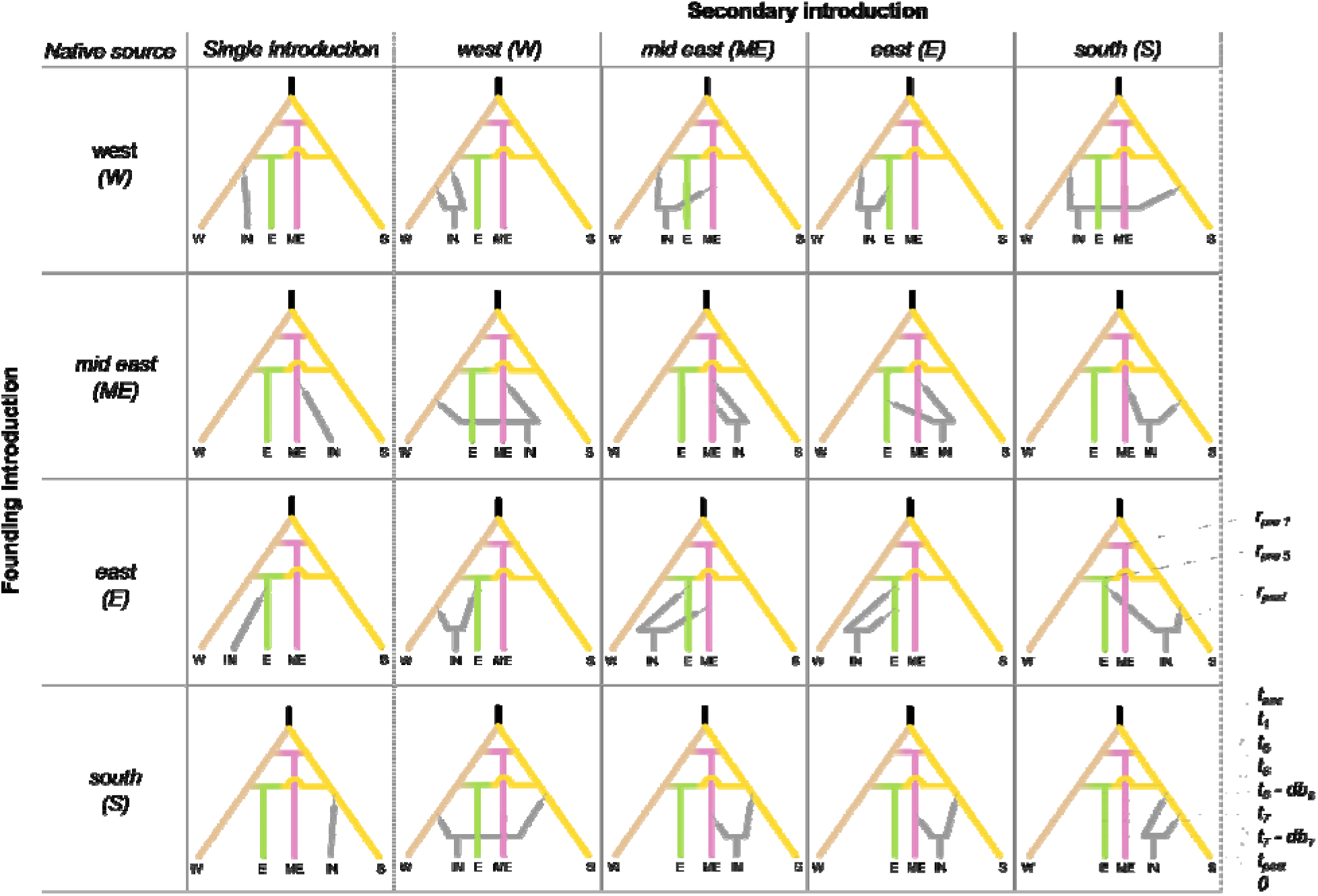
Graphical illustration of *Ambrosia artemisiifolia* introduction scenarios from divergent North American genetic units (W, E, ME, S) during initial (rows) and secondary (columns) introduction, tested using ABC-RF for European and Australian ranges (IN) independently. Thin lines indicate bottlenecks of duration *db_i_* with effective population sizes of *Nb_i_*. Time is not to scale. For detailed descriptions of parameters and prior intervals, refer to Table S13.

**Figure S5.**
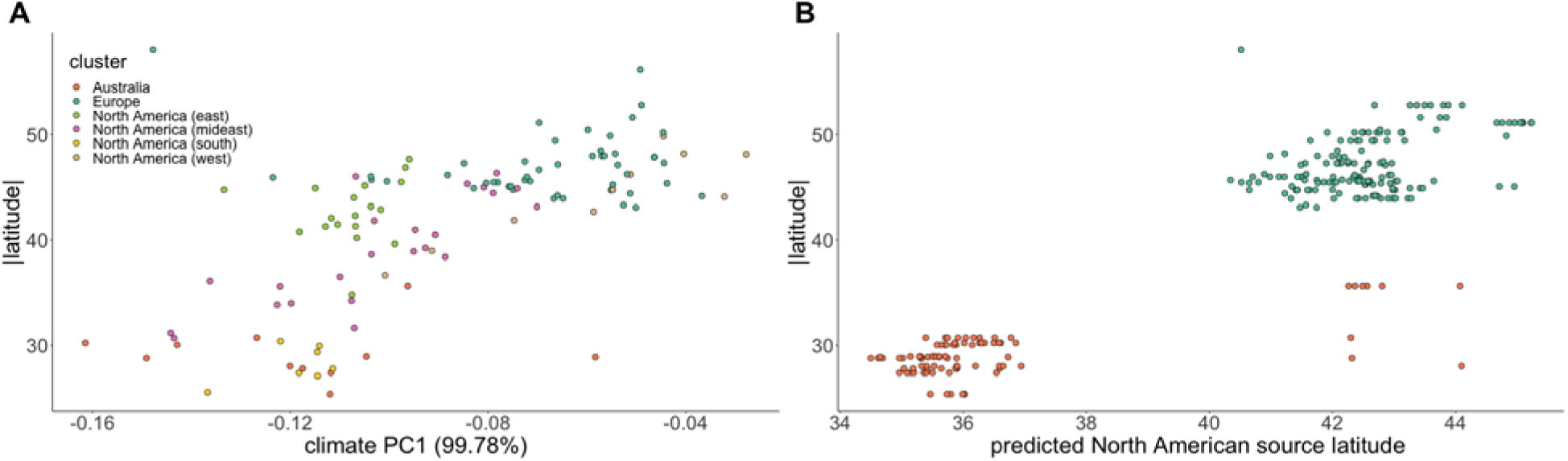
A. Relationships between population sampling locations in climate space (absolute latitude against the first principal component of four independent WorldClim variables [BIO1, BIO2, BIO12, BIO15]). **B.** Sampling location absolute latitude plotted against the *Locator*-predicted source location in North America for invasive-range samples.

**Figure S6.**
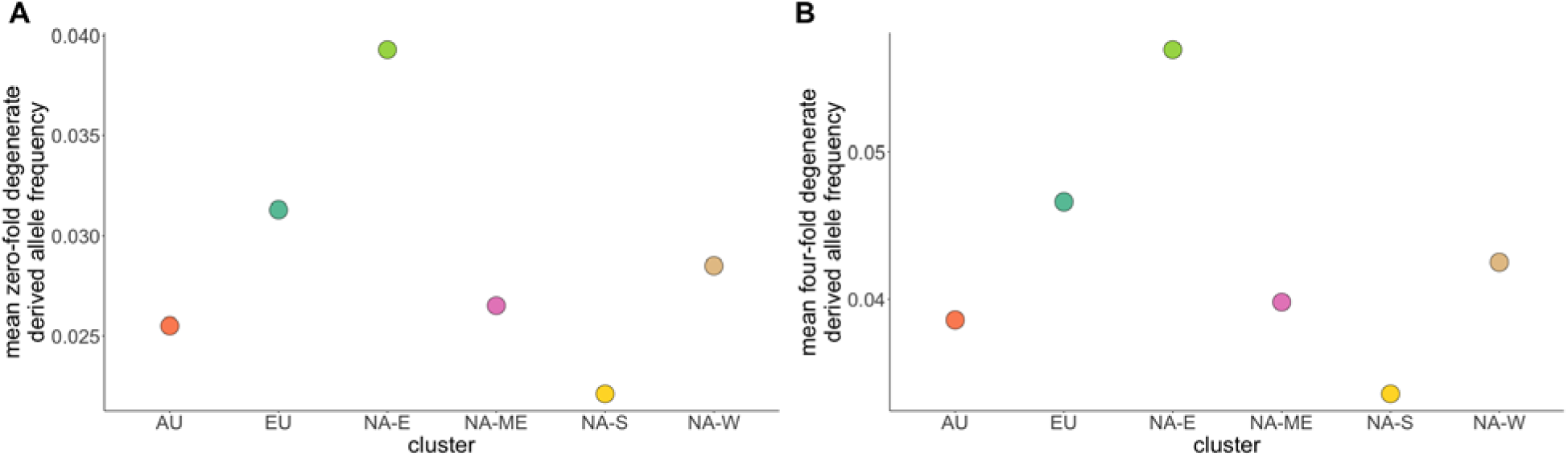
Mean allele frequency at zero-fold (**A**; as in Fig. 1F) and four-fold (**B**) degenerate sites for invasive ranges and North American clusters. Based on 95% CIs from bootstrapping sites, all points are significantly different from one another. CIs are too narrow to show on plots.

**Figure S7.**
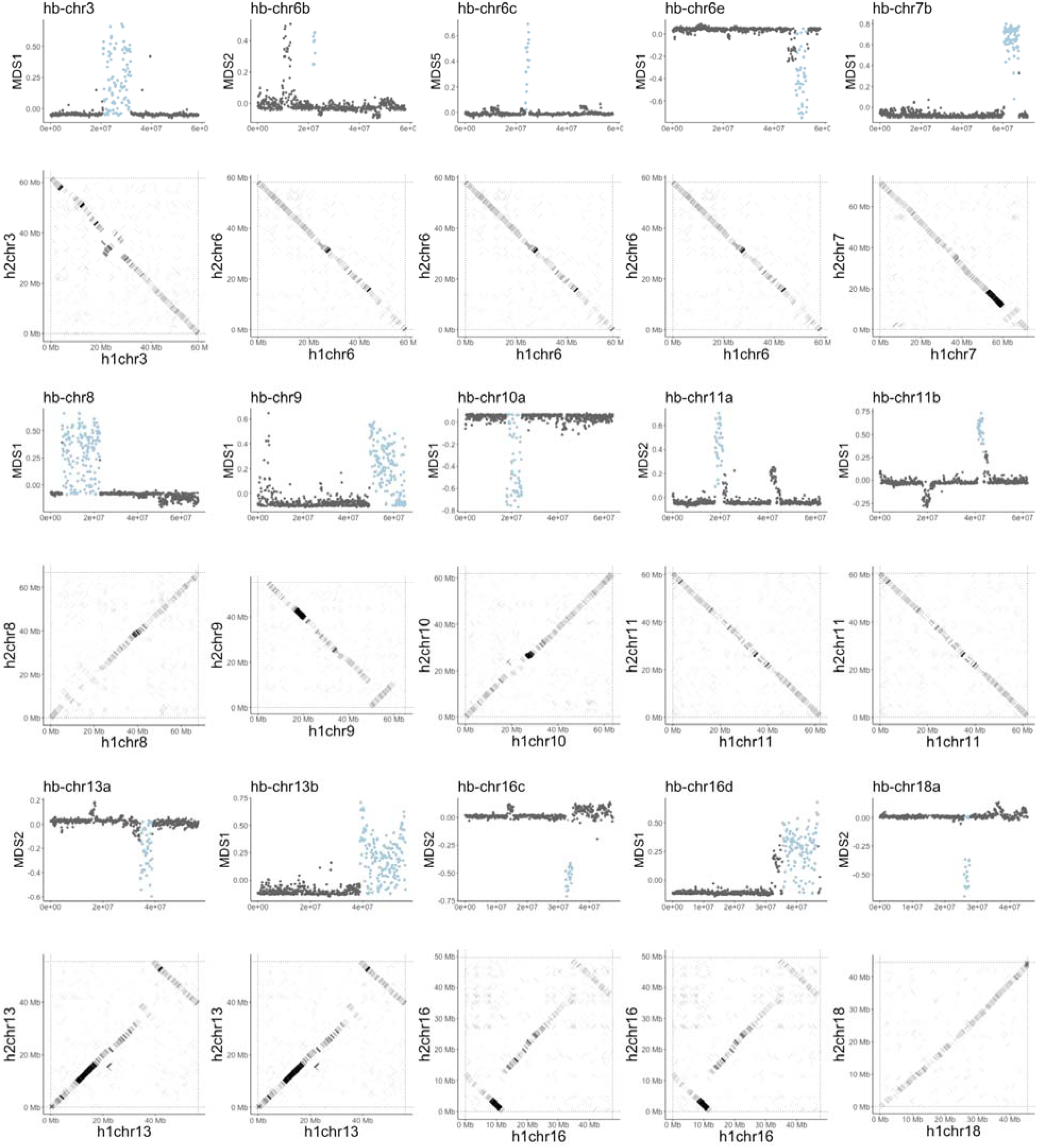
Divergent local population structure (MDS) in population-genomic data corresponds to inversion polymorphisms observed in alignments of homologous diploid reference chromosomes.

**Figure S8.**
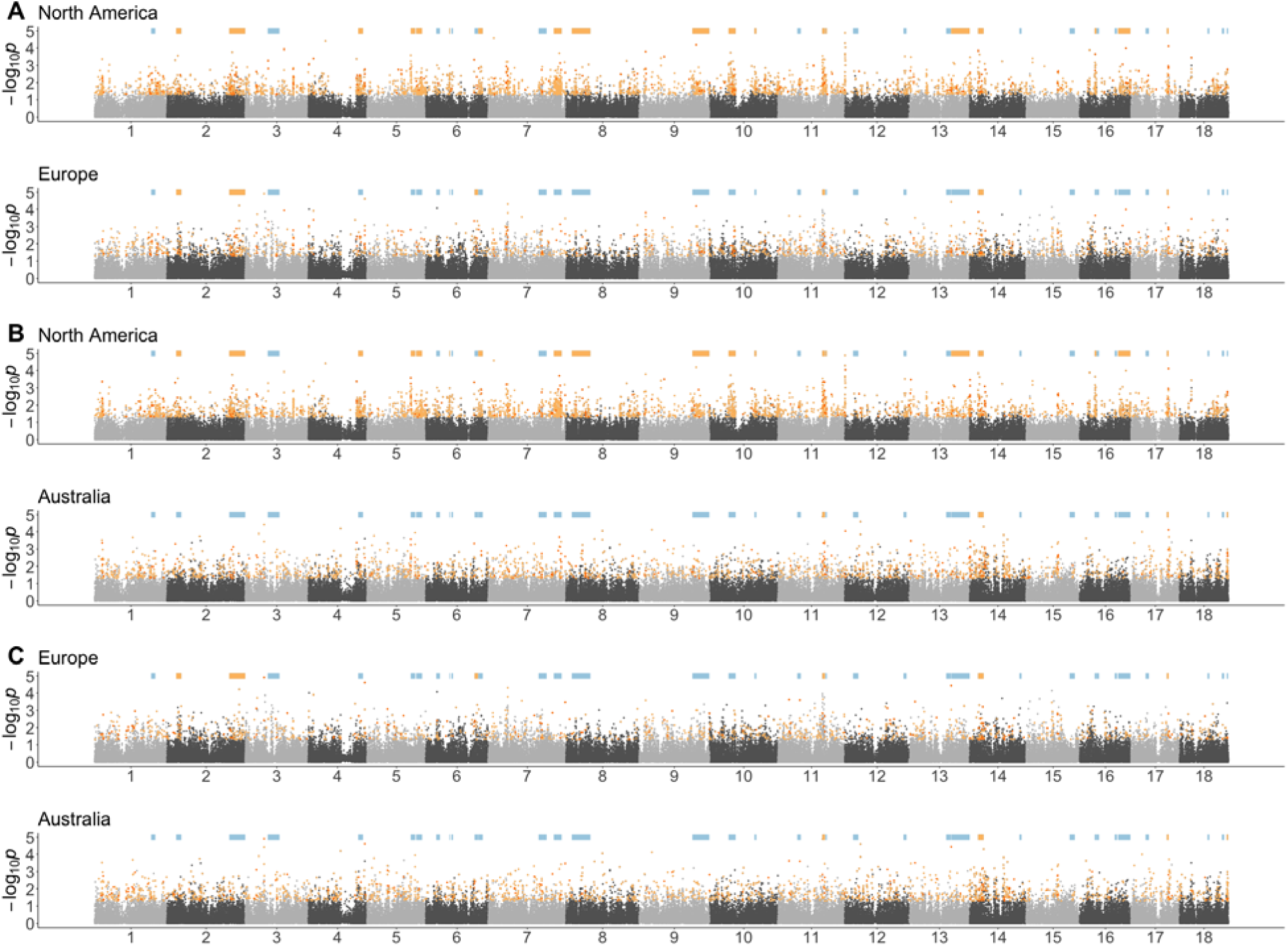
Manhattan plots (log-transformed empirical *p*-values for 10-kbp analysis window scores against genomic location) for XtX scans within each range separately. Orange highlights represent Xtx-EAA windows: the top 5% of XtX windows for each range that are also among the top 5% of EAA windows for at least one environmental variable in that range, with dark orange indicating outlier windows shared between each pairwise combination of ranges (**A**, **B** and **C**). Pale blue bars indicate the location o haploblocks. Orange bars denote haploblocks that are enriched for XtX-EAA windows.

**Figure S9.**
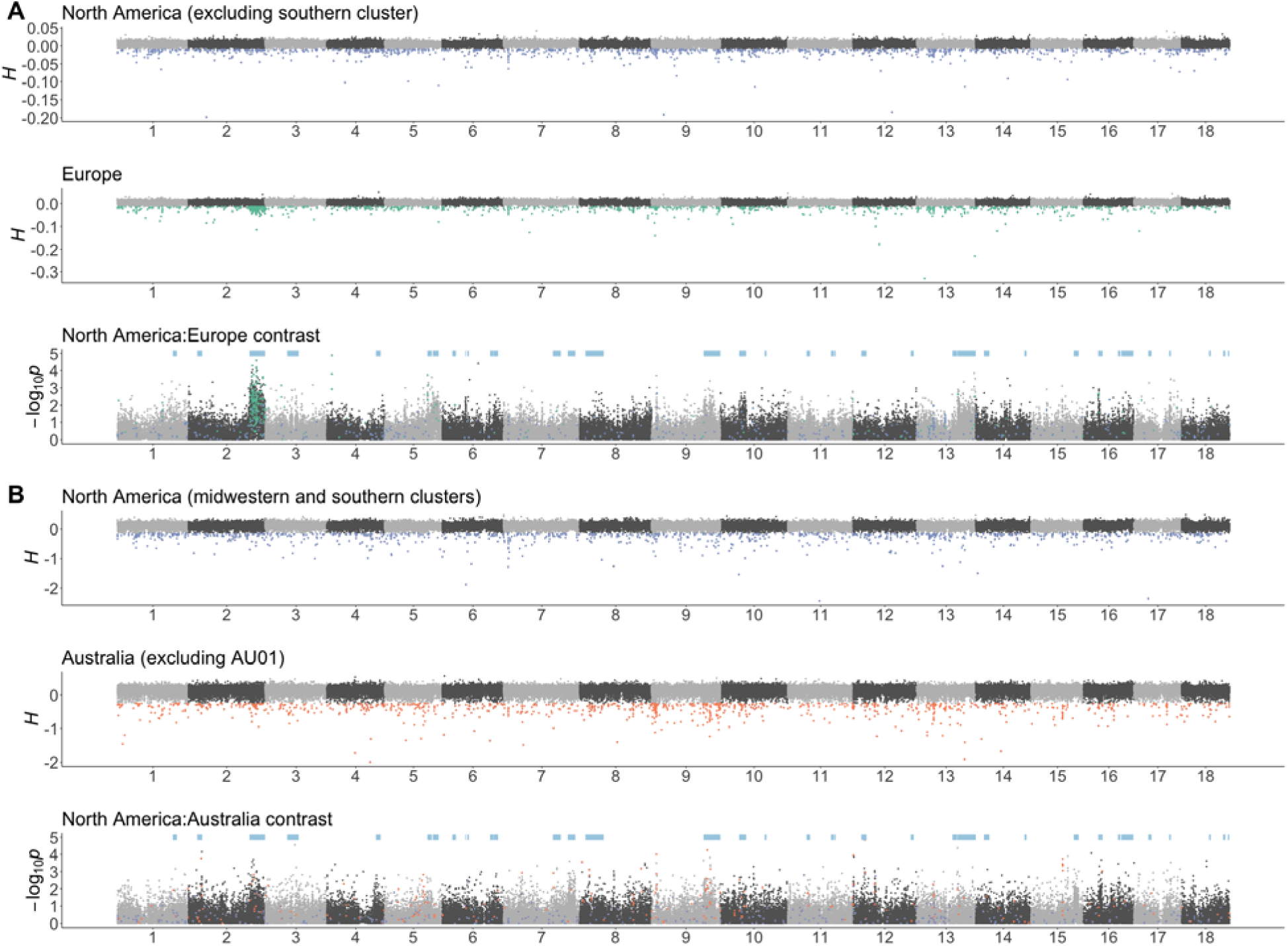
Manhattan plots for Fay and Wu’s *H* in individual ranges, and contrast scans between ranges for each native-invasive pair (**A** and **B**). 1% *H* outliers are coloured (North America: blue; Europe: green; Australia: red) in each *H* plot as well as the corresponding contrast plot. Pale blue bars indicate the location of haploblocks.

